# Can cryptic female choice prevent invasive hybridization in external fertilizing fish?

**DOI:** 10.1101/2021.12.17.473195

**Authors:** Tyler H. Lantiegne, Craig F. Purchase

## Abstract

Polyandrous mating systems result in females mating with multiple males. This includes the potential for unintended matings and subsequent sperm competition with hybridizing species, especially in the presence of alternative reproductive tactics (sneaker males). Cryptic female choice allows females to bias paternity towards preferred males under sperm competition and may include conspecific sperm preference when under hybridization threat. The potential becomes particularly important in context of invasive species that can novelly hybridize with natives. We provide the first examination of conspecific sperm preference in a system of three species with potential to hybridize: North American native Atlantic salmon (*Salmo salar*) and brook char (*Salvelinus fontinalis*), and invasive brown trout (*Salmo trutta*) from Europe. Using naturalized populations on the island of Newfoundland, we measured changes in sperm swimming performance, a known predictor of paternity, to determine the degree of upregulation to female cues related to conspecific sperm preference. Compared to water alone, female ovarian fluid in general had a pronounced effect and upregulated sperm motility (mean 53%) and swimming velocity (mean 30%). However, patterns in the degree of upregulation suggest there is no conspecific sperm preference in the North American populations. Furthermore, female cues from both native species tended to boost the sperm of invasive males more than their own. We conclude that cryptic female choice is too weak in this system to prevent invasive hybridization and is likely insufficient to promote or maintain reproductive isolation between the native species.

**Impact Summary:** Female mediated post-ejaculatory sexual selection, known as cryptic female choice, has only recently been researched in earnest, but has been documented across many taxa. This process allows females to bias paternity to favor a given male and can act as a filter to prevent fertilizations from unwanted males under sperm competition, including those of different species. In internal fertilizers like mammals, birds and insects, mechanisms of cryptic female choice can be very robust as the female can greatly modify the environment that sperm experience. In external fertilizers, females cannot control which males release sperm in close proximity to her eggs as she spawns with a chosen mate, but she can release reproductive fluids that act as a mechanism of cryptic female choice. In fishes, cryptic female choice is often mediated by ovarian fluid that is released with the eggs. This ovarian fluid alters sperm behavior, favoring certain males in situations of sperm competition. The mechanism is reportedly strong in native populations of European Atlantic salmon and brown trout, biasing paternity towards the female’s own species when eggs are under threat of hybridization under sperm competition. We examined cryptic female choice in three species of hybridizing salmonids on the North American island of Newfoundland, native Atlantic salmon and brook char, and invasive brown trout from Europe. Although the same species, salmon populations from both continents are quite distinct and our results suggest cryptic female choice is too weak in North American Atlantic salmon and brook char to prevent hybridization by invasive brown trout. We hope that this research inspires more work on cryptic female choice to better understand patterns across different species and locally adapted populations within species.

## Introduction

Sexual selection can occur via intrasex competition between individuals for access to mates and fertilizations, and mate choice for the opposite sex (Jones & Ratterman, 2009; Kuijper *et al*., 2012). Females are typically the choosier sex and select mates based on various attributes, including body odor (Ferkin, 2018) and courtship displays (Jennions & Petrie, 1997). Males, therefore, usually invest a large amount of energy into creating mating opportunities, while females invest comparatively more energy in the production of gametes and parental care (Bateman, 1948; Emery Thompson & Georgiev, 2014; Trivers, 1972). This difference of energetic expenditure between males and females creates situations where females may benefit from mating polyandrously, with more than one male, to better increase her chances of mating with high-quality males and producing good quality offspring (Firman, 2011).

In polyandrous mating systems, a female’s eggs are exposed to sperm from many males, creating sperm competition (Parker, 1970). Polyandry includes situations where females choose mates exhibiting dominant phenotypes (Morina *et al*., 2018), but other individual males circumvent female choice by resorting to alternative reproductive tactics to sneak fertilizations (Gross, 1996). In some systems, these tactics can result in fertilization by males of a different species, which facilitates hybridization (McGowan & Davidson, 1992; Tynkkynen *et al*., 2009; Garner & Neff, 2013). Across taxa, hybrid matings can result in highly variable outcomes, including speciation (Abbott *et al*., 2013), fertile or sterile hybrid offspring (Close & Bell, 1997), or no offspring due to failed fertilization, abortion, or abnormal development (Buss & Wright, 1958; Wilson *et al*., 1974; Chevassus, 1979). The potential for inviable or sterile offspring creates energetic waste (Remick, 1992); females have more to lose than males with each hybrid mating and thus should avoid hybrid fertilizations.

Under post-ejaculatory pre-zygotic sexual selection, females can bias sperm competition towards preferred males via cryptic female choice (Firman *et al*., 2017). The magnitude of this alteration between males can vary based on male relatedness (Landry *et al*., 2001; Yeates *et al*., 2009), perceived social status (Firman *et al*., 2017) and quality (Dean *et al*., 2011). Mechanisms of cryptic female choice in internal fertilizers include manipulating the duration of copulation, favouring males that provide greater stimulation during copulation, transferring favored sperm to better locations within the reproductive tract, discarding unwanted sperm, removing copulatory plugs, and changing internal conditions to be more or less favorable for sperm (Dixson, 2003; Eberhard, 2010; Pizzari & Birkhead, 2000). External fertilizers do not have this degree of control. Therefore, hybridization is more difficult to avoid in external fertilizers when unchosen males release sperm simultaneously with the female’s preferred mate.

However, externally fertilizing females can alter sperm behaviour using chemicals released with eggs, e.g., in mussels (Lymbery *et al*., 2017) and fish (Elofsson *et al*., 2006; Alonzo *et al*., 2016). Generally, these chemicals improve sperm swimming performance compared to a water-only environment (Lahnsteiner, 2002; Elofsson *et al*., 2006; Purchase & Rooke, 2020) and due to differential degree in response among males, subsequently bias fertilizations under sperm competition. Under hybrid matings, this form of cryptic female choice is known as conspecific sperm preference and allows a female to bias fertilization towards her own species (Yeates *et al*., 2013; Castillo & Moyle, 2019). In studies using paired species, conspecific sperm preference upregulates conspecific sperm swimming performance more than sperm from heterospecific males (Yeates *et al*., 2013; Castillo & Moyle, 2019), but how females relatively bias sperm performance across several species of potential fathers has not been investigated.

How does the strength of cryptic female choice via conspecific sperm preference vary with multiple species of differing degrees of phylogenetic relatability and/or likelihood of heterospecific matings? A good study system to examine this question is with external fertilizing salmonid fishes, as they are polyandrous (Haddeland *et al*., 2015), alternative reproductive tactics via sneak spawning is common (Esteve, 2005), and cryptic female choice mechanisms are reportedly strong (Butts *et al*., 2012; Yeates *et al*., 2013; Rosengrave *et al*., 2016) and readily manipulated. We chose three North American salmonids that can produce hybrids (Chevassus, 1979); native brook char (*Salvelinus fontinalis*) and Atlantic salmon (*Salmo salar*), and brown trout (*Salmo trutta*), which were introduced from Europe and are considered one of the top 100 worst invasive species in the world (Lowe *et al*., 2000). Brown trout create hybrids with both Atlantic salmon (Chevassus, 1979) and brook char (Buss & Wright, 1958), while native brook char and Atlantic salmon rarely produce viable offspring as a product of natural mating (Chevassus, 1979) – although actual mating rates between these species are not known and could be high. In their native Europe, salmon and trout are reported to show strong conspecific sperm preference that is mediated by ovarian fluid (Yeates *et al*., 2013), but North American populations of salmon and char have not been examined. Due to ovarian fluid creating a less hostile fertilization environment for sperm (Lahnsteiner, 2002; Elofsson *et al*., 2006; Lehnert *et al*., 2017), we hypothesized that (1) sperm respond positively to ovarian fluid when compared to swimming in only water, but (2) as a key mechanism to reduce the loss of eggs to hybridization, that ovarian fluid of all three species upregulates conspecific sperm more than heterospecific sperm, and (3) that there is a pattern in heterospecific upregulation that follows either taxonomic relationships or likelihood of spawning interactions occurring.

## Methods

### Experimental design

In the presence of sperm competition in natural matings, females are exposed to sperm from multiple males, which creates opportunities to bias paternity. To examine the potential for cryptic female choice, using a split-brood design, we took a sample of ovarian fluid from an individual female, split it into three aliquots, and exposed conspecific and two species of heterospecific sperm to it. We used a split-ejaculate design to quantify the sperm swimming performance of individual males in ovarian fluids and a water standard (Figure 1) and then determined their ratio. This ratio allowed us to quantify upregulation with a standardized value that is independent of differences between the values (e.g., an increase of 30 units from 30 to 60, or 40 units from 40 to 80, gives the same ratio) and thus controls for confounding variables such as individual differences in male sperm quality (Gage *et al*., 2004; Purchase & Moreau, 2012). A key prediction is that each ovarian fluid species upregulates conspecific sperm more than heterospecific sperm (Figure 1), ie all ovarian fluid species should create standardized values >1 for all sperm species, but the ratio should be highest for conspecific sperm.

**Figure 1:**
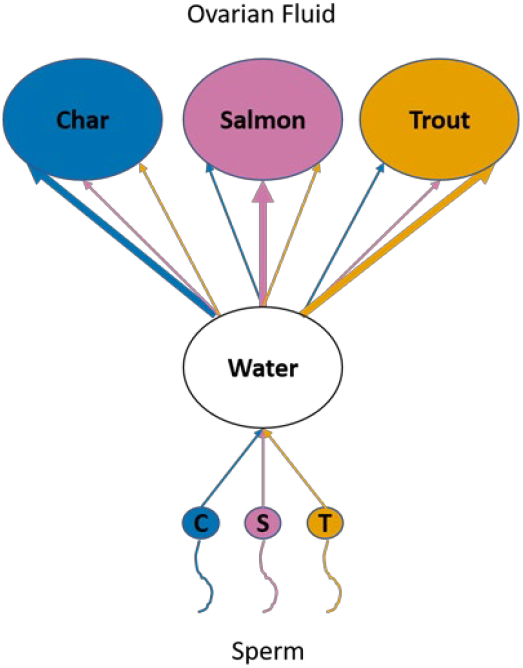
Schematic of the conceptual design. Small circles with tails are sperm, while large circles are ovarian fluid and water. Arrows represent sperm velocity in water or ovarian fluid (the same semen sample was tested in both as separate aliquots of individual sperm). Ovarian fluids were predicted to show conspecific sperm preference, indicated by greater upregulation of conspecific sperm (bolded arrows) over heterospecific sperm (un-bolded arrows). Upregulation was quantified as the ratio of sperm swimming performance in ovarian fluid compared to that in water, which controls for individual variation in male quality (variable performance among males in water). Experimental replication was achieved with 12 groups of fish, and sperm activations were technically repeated 3 times for each comparison.

Sperm swimming performance comparisons were conducted over a series of experimental replicates (unique groups of fish), each containing one female and one male of each species. In each of these replicates, samples of ovarian fluid and sperm from each of our three study species were exposed to one another. Each sperm activation was technically repeated three times. Every experimental replicate tested each male in water, as well as individual samples of the three ovarian fluids (3 males per replicate, each male’s sperm activated 12 times, for a total of 36 sperm swimming comparisons per replicate). We ran 12 experimental replicates with different fish, with two replicates occurring on a given day. In two of the total 36 sampled males, preliminary assessment of semen quality was very poor, and we replaced that fish with the male from the other replicate on that day. Over the course of the study, we used 70 fish: 12 females of each species, and 12 brown trout, 11 brook char, and 11 Atlantic salmon males. To simplify analyses, we subsequently treated the two reused males as independent, as they were used with different females. We produced 432 sperm swimming comparisons (12 experimental replicates * 3 species of male * 4 sperm activation solutions * 3 technical replicates).

### Fish collection

Fish handling was conducted under approval of Memorial University’s Animal Care Committee (permit 20190378). Population sizes of our study species were large enough not to be negatively affected by sampling. Fish were sourced from different waterbodies, but to avoid potential confounding variables, care was taken to ensure that all sperm could be examined at “exactly” the same amount of time from collection. As a species, salmon co-evolved with trout and char, but North American salmon have been isolated from European salmon for 600,000 years (Lehnert *et al*., 2020), and are genetically different (e.g. Hartley, 1987). Brown trout co-evolved with salmon in Europe but were not exposed to brook char, while our North American salmon co-evolved with brook char. Details of each sampled population are described below.

Wild native Atlantic salmon were sourced from the Exploits River in Newfoundland, Canada (48.93 N, 55.67 W). These co-occur with native brook char in this watershed. Fish were trapped in the fishway on Grand Falls on September 7, 2018 and transferred to tanks on September 30, following previous protocols (Rooke *et al*., 2019). At ∼11 AM on gamete collection days from November 2 – 14, individuals were anesthetized with MS-222, paper toweled dry, and then stripped of gametes via ventral massage. Semen was stripped into plastic bags and eggs into glass jars.

Wild native brook char were collected from Star Lake in Newfoundland, Canada (48.58 N, 57.23 W). This is part of the Exploits River watershed, but there are no reports of salmon occurring within this particular lake. Char were captured via fyke net from September 21 to October 5, 2018, transported via truck, and housed in tanks at the same facility as the salmon. Brook char were fed a diet of mealworms until October 5 and then fed 4mm biobrood pellets for the remaining duration of captivity (salmon do not eat before spawning and were thus not fed). Brook char were anesthetized with MS-222. Females were stripped over the last week of October, the eggs were filtered out – see below, and ovarian fluid frozen (Purchase & Rooke, 2020). Brook char males were stripped of semen immediately (minutes) after the salmon. Fresh char semen and frozen ovarian fluid were stored in 1.5 ml Eppendorf tubes. Gametes from salmon and char were transported on ice and received at the laboratory in St. John’s at ∼11 PM; all experimental procedures were done overnight and completed within 24 hours of gamete collection.

Brown trout have not yet invaded the Exploits River watershed from which char and salmon were collected in central Newfoundland (Westley & Fleming, 2011). Trout were introduced from Scotland in the late 19^th^ century (Hustins, 2007) into watersheds surrounding St. John’s and have since invaded throughout southeastern Newfoundland. As invaders, these fish have been documented to hybridize with (McGowan & Davidson, 1992) and outcompete (Sorensen *et al*., 1995) native salmonids. Based on a generation time of 3-5 years, there were 27-35 generations of brown trout in Newfoundland at the time of collection. Wild, non-native brown trout used in this study were captured via dipnet in tributaries of Windsor Lake (47.60 N, 52.78 W), in St. John’s Newfoundland, where there are brook char but no Atlantic salmon. Trout were anesthetized immediately after capture, measured for length, fin-clipped to avoid double sampling on different days, and stripped for gametes into plastic containers. Through coordinated field activities, trout stripping took place on the same days and at the same time (<1 hour) as Atlantic salmon and brook char stripping. Trout gametes were kept on ice for ∼12 hours, the same duration as char and salmon before use. Animal care protocols did not allow the use of MS-222 with fish (trout) being released back into the wild, and thus clove oil was used. Both anesthetics have been shown to have no significant effects to gametes when used prior to gamete collection (Holcomb *et al*., 2004).

### Gamete preparation

An aliquot of semen (0.5 ml) from each male was centrifuged at 4100 g for 10 mins at 5°C to separate seminal fluid from sperm. This seminal fluid acted as a non-activating diluting agent to decrease the density of other aliquots from the same fish (Purchase & Moreau, 2012) at a 1:75 sperm to seminal fluid ratio. This minimized sperm clumping and allowed for high-quality sperm data measurement. The ovarian fluid was filtered from eggs with a fine-mesh aquarium net and refrigerated at 4°C in a glass beaker. Ovarian fluid activating solutions were made at 33% concentration with water. Bovine serum albumin was included in the sperm activating solution at a concentration of 1:1000 to prevent sperm from adhering to the microscope slide (Beirão et al., 2014; Beirão et al., 2015).

Sperm swimming performance was recorded using a Prosilica GE680 camera attached to an inverted Leica DM IL LED microscope, with a 20× phase contrast objective. Approximately 1µl of diluted semen was put on the edge of the chamber of a Cytonix 2 chambered slide, which had been cooled to ∼ 9°C with a custom Physitemp TS-4 system. The semen was then flushed into the chamber by 395 µl of the sperm activating solution (the test treatment). This activated the sperm and marked the start of the video, which was taken at 80 fps. The first 6s post-activation were used to locate an area of suitable sperm density and focus the microscope. Videos were captured using Streampix software, and quality checked for sperm density, motility, and proper microscope focus before being accepted into the data pool. If a video was deemed poor quality, the entire sperm activation process was repeated until three adequate videos were attained for technical replication (see above). Data from these three videos were averaged before analyses.

### Data analyses

Sperm swimming performance was determined from 6.0 to 20.0s post-activation, using the Computer Assisted Sperm Analysis (CASA) plugin in ImageJ with a tracking interval of 0.5s (Purchase & Earle, 2012), Supplemental Table 1. Two sperm swimming performance traits were used in analyses, as these have been shown to be related to paternity under sperm competition (Gage *et al*., 2004; Evans *et al*., 2013; Young *et al*., 2013; Alonzo *et al*., 2016; Lehnert *et al*., 2017); the percent of the sperm cells within an ejaculate that are motile (MOT), and of the motile cells, their curvilinear swimming velocity (VCL). How ovarian fluid modified these parameters in different species of sperm – controlling for individual variation in sperm quality using a water standard, was our metric for determining conspecific sperm preference and thus the ability of females to exert cryptic female choice. To simplify analyses, we elected to focus this comparison to the earliest sperm post-activation time period available (6.0-6.5 seconds) as fertilizations happen quickly (Hoysak & Liley, 2001; Rosengrave *et al*., 2016; Beirão *et al*., 2019), and thus represents the most important time interval for females to modify. Changes over the full recorded time of sperm swimming are presented in the Supplement.

Assumptions of parametric statistics were tested by examining model residuals. To test hypothesis (1) that sperm respond positively to ovarian fluid when compared to swimming in only water, we constructed mixed effects generalized linear models for motility (binomial error) and VCL (normal error), Equation 1. Both models used the fixed effect of sperm activating solution (water or ovarian fluid – the average of all types) and the random effect of male ID as the independent variables.

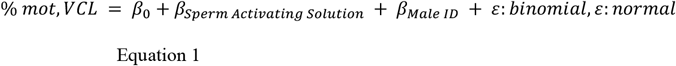

To evaluate hypothesis (2) that ovarian fluid upregulates conspecific sperm more than heterospecific sperm we used two approaches. Both velocity and motility (as standardized motility values were no longer proportions) were tested with normal distributed error. First, we broke ovarian fluids into two categories for each male (conspecific {1 female} or heterospecific {average of 2 females}). We used standardized swimming performance (the ratio in ovarian fluid to water) as the dependent variable, ovarian fluid type (conspecific or heterospecific) as a fixed independent variable, and male ID as a random independent variable (Equation 2). A significant result from this model would indicate that across 36 females (ignoring their species) on average, upregulation of conspecific sperm would be different than the average of two species of heterospecific sperm.

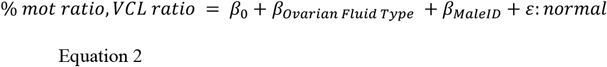

Second, to conduct analyses at a finer resolution in case only some species of ovarian fluid support conspecific sperm preference, and to test hypothesis (3) that there are patterns of upregulation among the two heterospecific species of sperm within each ovarian fluid, we constructed a generalized mixed-effects model and used standardized motility and velocity as dependent variables, and sperm species, ovarian fluid species (char, salmon, or trout), and their interaction as fixed independent variables, and male and female ID as random independent variables (Equation 3). If the interaction was significant, the model was broken down by ovarian fluid species, and if necessary analyzed post-hoc with a Tukey test to determine differences among male species in each ovarian fluid.

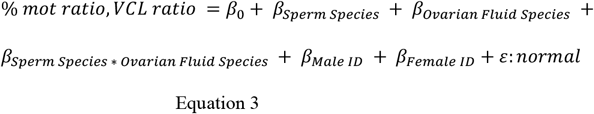

## Results

Despite individual variation among males in sperm quality, on average sperm of all three species had similar swimming characteristics that declined rapidly post-activation (Figure 2). A declining function was expected, so we simplified subsequent results and focused on the most biologically relevant time for sperm competition; the earliest we could capture, 6s. Our first hypothesis, that sperm respond positively to ovarian fluid when compared to swimming in only water, was supported (Figure 3 positive slopes, Figure 4 ratios >1). Individual male performance was visualized as a reaction norm following Purchase et al. (2010) to show this upregulation between water and all ovarian fluids combined (Figure 3). Motility (df=1, F=6.15, p=0.018) and curvilinear velocity (df=1, F=83.67, p<0.001) were both significantly upregulated at 6s on average by 53% (ratio of OF/W = 1.53) and 30% (ratio of OF/W = 1.30), respectively. Using this approach, we were then able to use the unique ratios of upregulation by each ovarian fluid for each species of sperm to investigate patterns.

**Figure 2:**
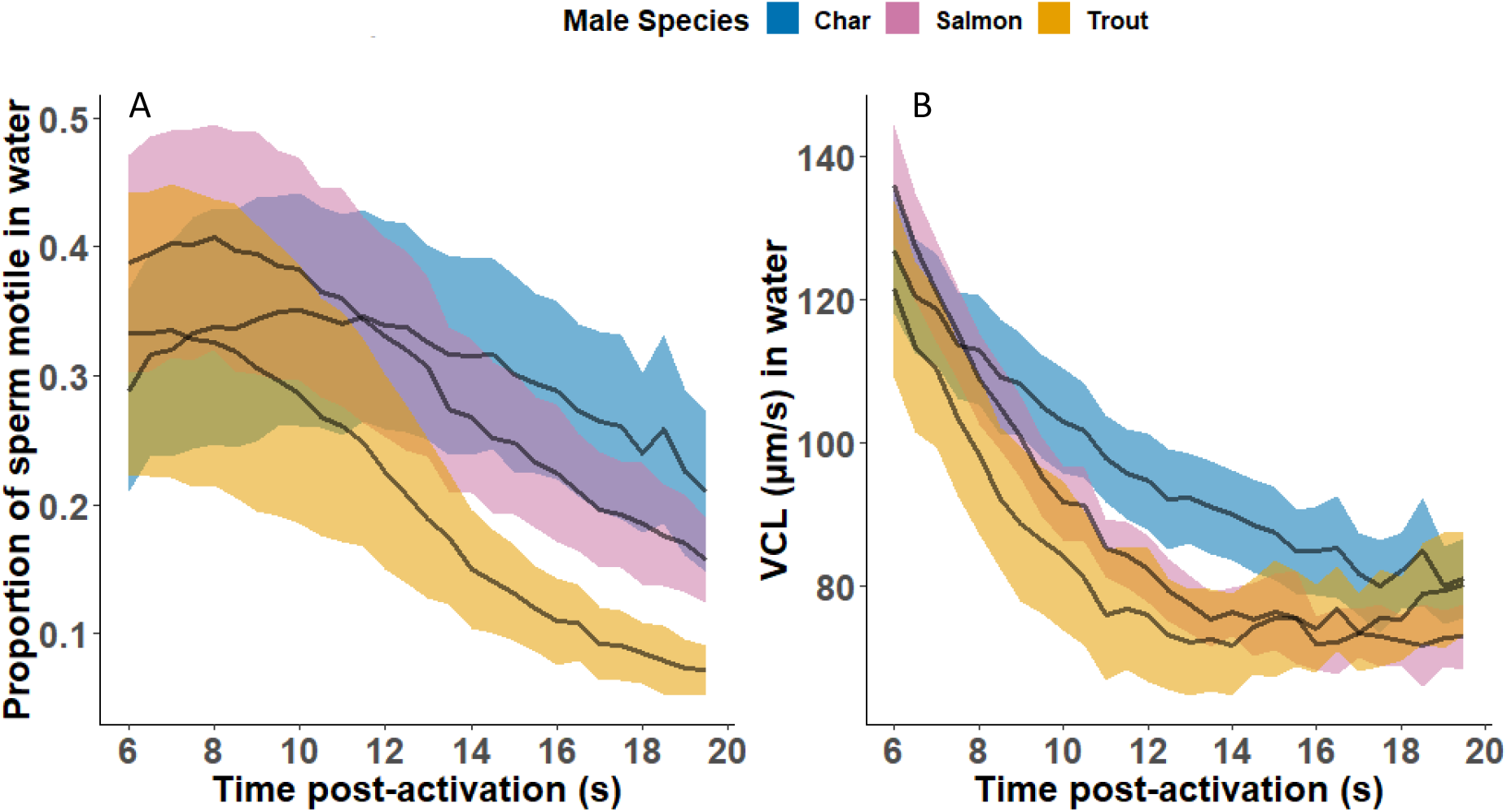
Sperm swimming performance in water (panel A: proportion motile, B: curvilinear swimming velocity [VCL]) from 6.0-20.0s post-activation. Black lines represent the average at each 0.5s interval, and colored bands are 2*standard error among 12 individual males within a species (blue = char, pink = salmon, orange = trout).

**Figure 3:**
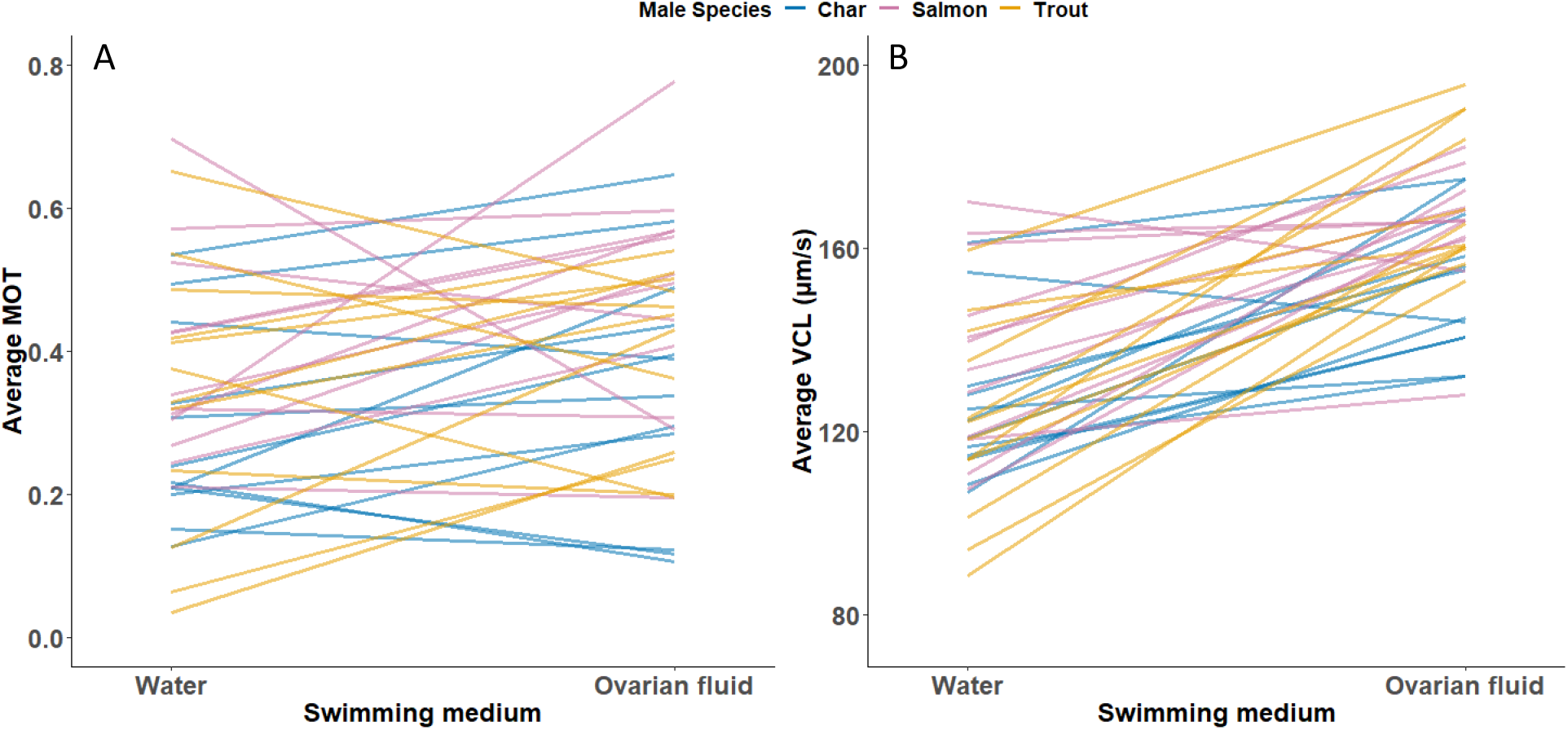
Reaction norms (A= proportion sperm motile, B= curvilinear swimming velocity) comparing sperm swimming performance from 6.0-6.5s post-activation in water to the average value in 3 ovarian fluid species. Each line represents an individual male (blue=char, pink=salmon, orange=trout) and is created by two points; means for water represent three technical replicate activations for each male, while those for ovarian fluid are from nine activations (3 technical replications from each of 3 species of ovarian fluid). Positive slopes indicate up-regulation of sperm swimming by ovarian fluid. Standardized (ovarian fluid / water) ratios >1.0 indicate positive up-regulation and were on average 1.53 for MOT and 1.30 for VCL.

**Figure 4:**
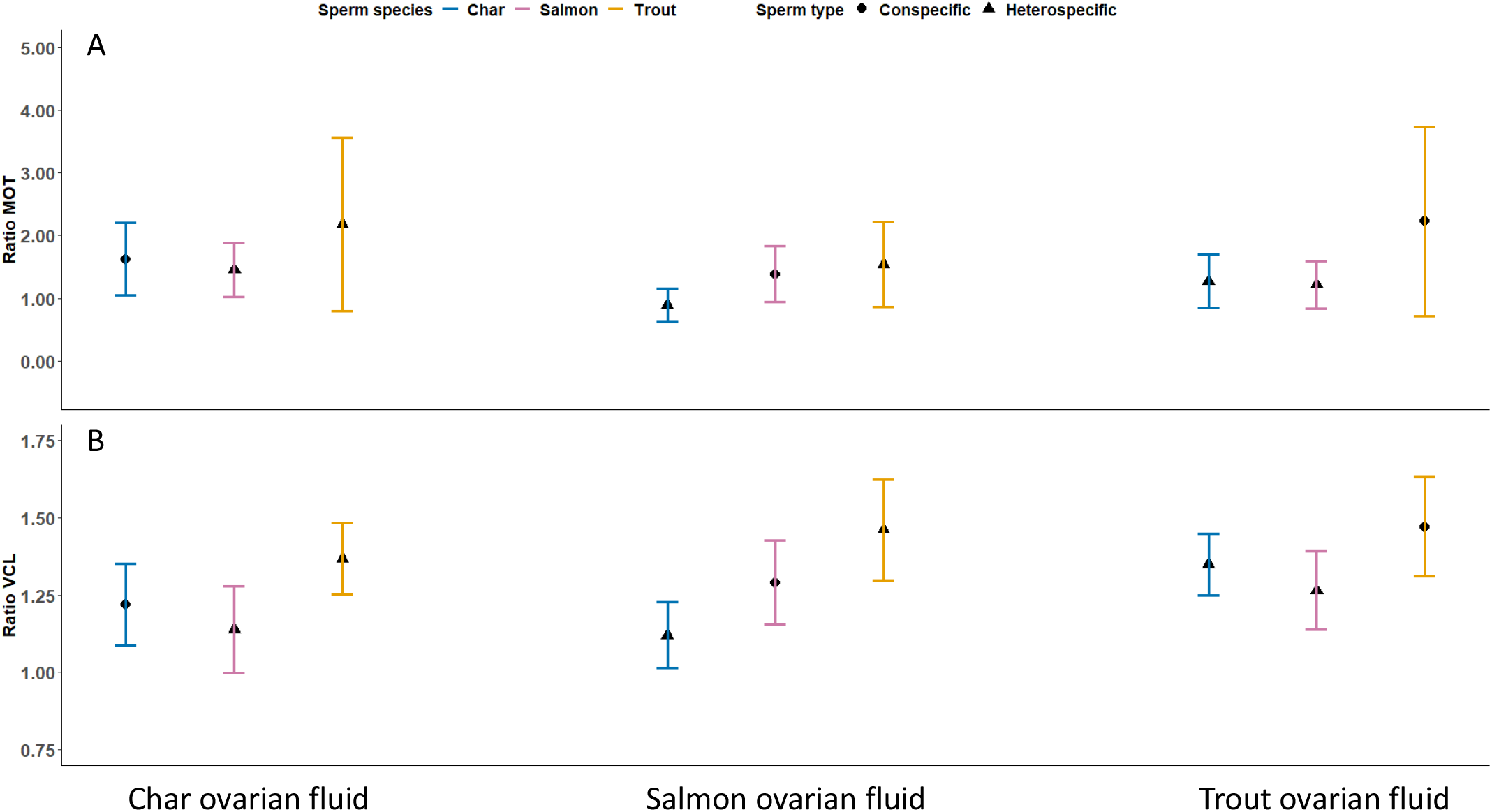
Ratio of (A) sperm motility (% motile) and (B) curvilinear velocity (VCL µm/s) in specific ovarian fluid compared to water from 6.0-6.5s post-activation – any value above 1.0 indicates ovarian fluid up-regulated sperm swimming performance. Black shapes represent the average (circles are conspecific sperm to the ovarian fluid, triangles are heterospecific sperm), and colored brackets 2*standard error among 12 males and females within a species.

Our second hypotheses, that across species, ovarian fluid consistently changes sperm swimming to enable conspecific sperm preference was not supported by either of our approaches (equations 2 and 3). There was variation in how much ovarian fluid upregulated sperm by male species, but trends were not consistent at 6s (Figure 4) or throughout the full recorded sperm swimming period (Supplementary Figure). At 6s post-activation (Figure 4 - circles vs triangles), there was no statistically significant difference in how much ovarian fluid upregulated conspecific vs. heterospecific sperm (two heterospecific species averaged – equation 2) for sperm motility (Figure 4a; df=1, F=1.37, p=0.250) or velocity (Figure 4b; df=1, F=2.36, p=0.133).

For our second approach to hypothesis 2 (higher resolution – equation 3), and hypothesis 3, that there is a consistent pattern in how ovarian fluid upregulates sperm of different heterospecific species, we examined effects at the species level. Reporting general trends (Figure 4), trout ovarian fluid did weakly (see below) upregulate trout sperm more than heterospecific sperm (indicative of conspecific sperm preference), but char and salmon ovarian fluid tended to upregulate trout sperm more than sperm of their own species (not supporting conspecific sperm preference) – note trout sperm swimming performance in water was not higher than the other species (Figure 2). For motility (Figure 4a) the different ovarian fluid species did not upregulate sperm species differently (interaction not significant; df=4, F=0.69, p=0.601), indicating no pattern of upregulation to support hypotheses 2 or 3. The interaction between male and female species was significant for velocity (df=4, F=6.20, p=0.004). The VCL model was broken down and analyzed separately for each ovarian fluid species. There were no differences in upregulation among the 3 sperm species by char ovarian fluid (df=2, F=3.20, p=0.532) or trout ovarian fluid (df=2, F=4.49, p=0.098) – providing no support for hypotheses 2 or 3. There was a difference in sperm upregulation by salmon ovarian fluid (df=2, F=6.17, p=0.005), however not in a manner to support hypothesis 2 (Figure 4b), as trout sperm were upregulated significantly more than char sperm (Tukey: df=33, t=-3.51, p=0.004), but there was no difference between either trout and salmon (Tukey: df=33, t=-1.57, p=0.200) or char and salmon sperm (Tukey: df=33, t=-1.56, p=0.200).

## Discussion

Conspecific sperm preference is the last line of defence against hybridization of a female’s eggs and has been demonstrated in taxa as diverse as mussels (Klibansky & McCartney, 2014), crickets (Howard *et al*., 1998; Tyler *et al*., 2013), birds (Pizzari & Birkhead, 2000; Wagner *et al*., 2004), and European populations of salmon and trout (Yeates *et al*., 2013). We therefore expected ovarian fluid mediated upregulation (documented to enable conspecific sperm preference) of sperm swimming performance would be strong in our hybridizing salmonids. However, while ovarian fluid consistently upregulated sperm swimming performance when compared to water, it did not do so more for conspecific than heterospecific sperm, and thus these females cannot bias paternity towards their own species in this way. Given it is the only known mechanism possible for external fertilizing fish, we therefore conclude that cryptic female choice is too weak to promote or maintain reproductive isolation between native North American Atlantic salmon and brook char, nor can it reduce hybridization by invading brown trout.

Ovarian fluid clearly upregulated sperm motility and velocity. Other studies have shown that components of ovarian fluid (Rosengrave *et al*., 2009; Lehnert *et al*., 2017) prolong sperm lifespan and increase sperm velocity in these and related taxa (Urbach *et al*., 2005; Elofsson *et al*., 2006; Evans *et al*., 2013; Purchase & Rooke, 2020). Since this function of ovarian fluid was strongly demonstrated, we expected to see conspecific sperm preference. However, ovarian fluid did not upregulate conspecific sperm more than heterospecific sperm in Newfoundland salmonids. Trout ovarian fluid did weakly upregulate trout sperm more, but trout sperm also tended to be upregulated more (not significantly) than the others in the two heterospecific ovarian fluids. This result is surprising because of the high cost to females from fertilization by heterospecific males. Atlantic salmon and brook char do not create viable adult hybrids (Chevassus, 1979) – but hybrid mating rates are unquantified and could be high, brown trout and brook char create sterile adults – known as tiger trout (Buss & Wright, 1958), and Atlantic salmon and brown trout create sterile F2s (Chevassus, 1979). In all cases, hybrid fertilizations of a female’s eggs create evolutionary dead ends. Preventing hybrid fertilizations under heterospecific sperm competition would therefore be highly adaptive.

Yeates et al. (2013) examined hybridization with European populations of Atlantic salmon and brown trout. They found that ovarian fluid was strongly linked to conspecific sperm preference and that the egg itself did not have any protections against hybridization. Ours is the first investigation of conspecific sperm preference for any brook char population. Our study system also has Atlantic salmon that have been isolated from European salmon for 600,000 years (Lehnert et al., 2020), introduced (<150 years) and invasive brown trout, and documented wild hybridization (McGowan & Davidson, 1992). Although as species, salmon and trout evolved together, it is possible allopatric populations of North American salmon may have lost the ability of their European cousins to prevent hybridization by brown trout in sperm competition. Our results suggest cryptic female choice via conspecific sperm preference in upregulation of sperm swimming performance is too weak to prevent hybridization in our study populations. However, to more thoroughly examine cryptic female choice as a means to prevent hybridization in salmonids, more species, and populations over a range of allopatric and sympatric distributions should be investigated. Sperm swimming performance evaluations should also be followed by sperm competition experiments to confirm effects on paternity.

## Acknowledgements

Funding was provided via Memorial University, and grants to CP from the Atlantic Salmon Conservation Foundation, the Natural Sciences and Engineering Research Council of Canada, the Canada Foundation for Innovation, and the Research and Development Corporation of Newfoundland and Labrador. We thank the staff of the Environmental Resources Management Association, particularly Terry Paul and Darren Ryan, for the acquisition and shipping of salmon and char gametes. Assistance in collecting trout gametes and conducting sperm swimming comparisons was provided by Terry Sullivan, Madison Philipp, Coady Fitzpatrick, Sydney London, Taylor Hughes, and Alexander Flynn. We thank Ian Jones, Travis Van Leeuwen, Peter Westley and Simone Immler for providing comments on an earlier version of the MSc thesis that contained this manuscript. The authors declare no conflicts of interest.

## Author Contributions

CP conceived the study. TL and CP designed the experiment. TL collected and analyzed the data, with input from CP. TL and CP co-wrote the manuscript.

---

**Supplemental table 1:** Casa parameters for sperm swimming characteristic analysis

| CASA Parameter | Value |
| --- | --- |
| Minimum sperm size (pixels) | 40 |
| Maximum sperm size (pixels) | 440 |
| Minimum track length (frames) | 97 |
| Maximum sperm velocity between frames (pixels) | 8 |
| Minimum VSL for motile ($\mu\text{m/s}$) | 3 |
| Minimum VAP for motile ($\mu\text{m/s}$) | 20 |
| Minimum VCL for motile ($\mu\text{m/s}$) | 45 |
| Low VAP speed ($\mu\text{m/s}$) | 5 |
| Maximum percentage of path with zero VAP | 1 |
| Maximum percentage of path with low VAP | 25 |
| Low VAP speed 2 ($\mu\text{m/s}$) | 25 |
| Low VCL speed ($\mu\text{m/s}$) | 35 |
| High WOB (percent VAP/VCL) | 80 |
| High LIN (percent VSL/VAP) | 80 |
| High WOB two (VAP/VCL) | 50 |
| High LIN two (VSL/VAP) | 60 |
| Frame Rate (frames per second) | 80 |
| Microns per 1000 pixels | 1075 |

**Supplemental figure 1:**
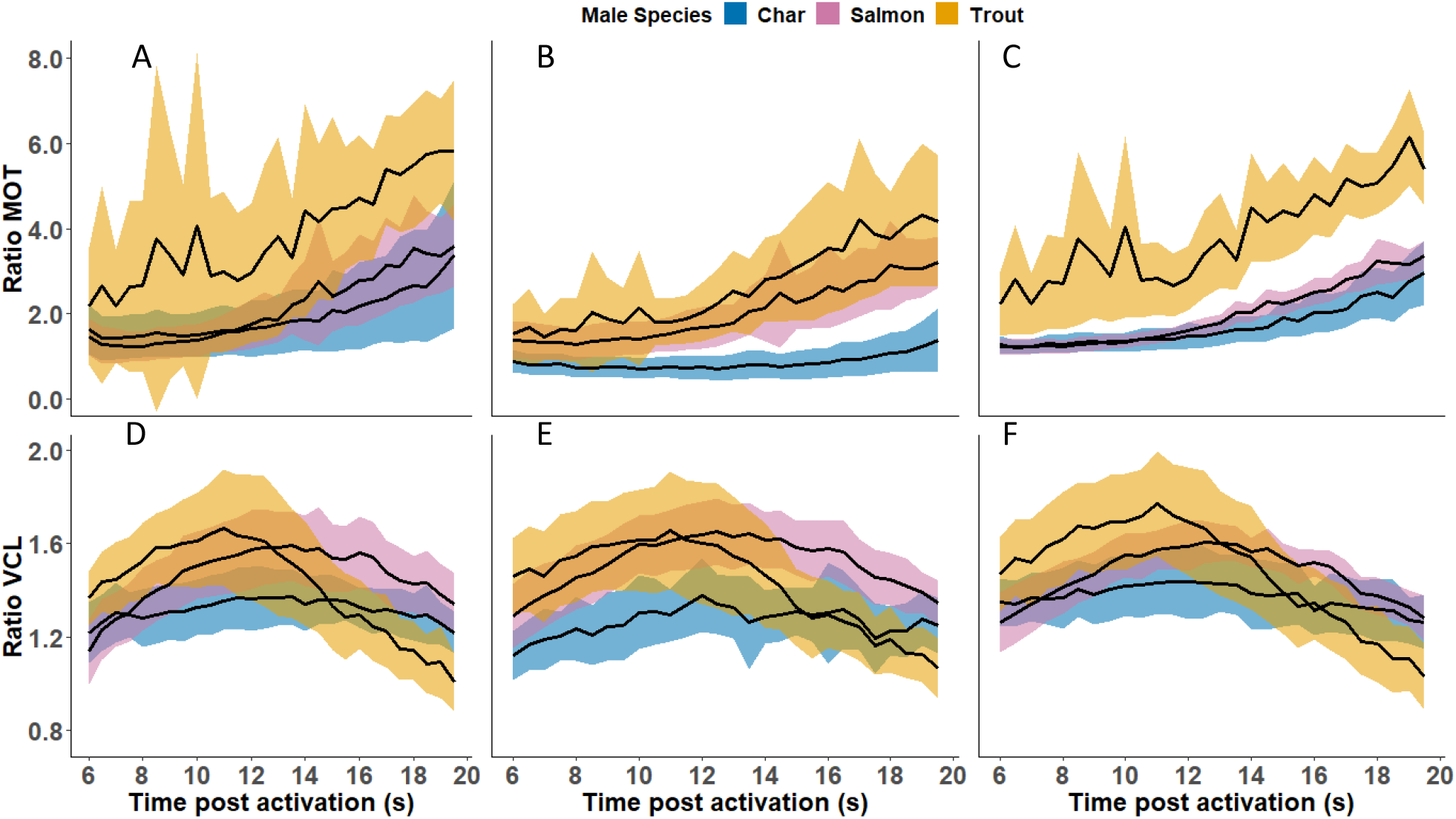
Ratio of char (blue), salmon (pink), trout (orange) sperm motility and curvilinear velocity (VCL µm/s) in ovarian fluid compared to water from 6 to 20s post activation. Black lines represent the average at each 0.5s and colored bands represent the 2*standard error for the 12 males and females within a species. Panels A B C show the ratio of sperm motility in water over char, salmon, and trout ovarian fluid, respectively. Panels D E F show the ratio of VCL in char, salmon, and trout, respectively.

